# Efficient detection of transposable element insertion polymorphisms between genomes using short-read sequencing data

**DOI:** 10.1101/2020.06.09.142331

**Authors:** P. Baduel, L. Quadrana, V. Colot

## Abstract

Transposable elements (TEs) are powerful generators of major-effect mutations, most of which are deleterious at the species level and maintained at very low frequencies within populations. As reference genomes can only capture a minor fraction of such variants, methods were developed to detect TE insertion polymorphisms (TIPs) in non-reference genomes from short-read sequencing data, which are becoming increasingly available. We present here a bioinformatic framework combining an improved version of the SPLITREADER and TEPID pipelines to detect non-reference TE presence and reference TE absence variants, respectively. We benchmark our method on ten non-reference *Arabidopsis thaliana* genomes and demonstrate its high specificity and sensitivity in the detection of TIPs between genomes.

## 1 Introduction

Transposable elements (TEs) have the ability to move across the genome and as such they constitute powerful endogenous mutagens. The major-effect mutations they generate when they insert near or within genes are mostly deleterious and such variants are therefore submitted to strong purifying selection^1^. Consequently, the vast majority of TE insertions with phenotypic impact segregate at very low frequency across populations. Yet, most genome-wide studies of TEs are limited to those TE sequences that are annotated in reference genomes and thus cannot properly assess the contribution of TE mobilization to the creation of genetic novelties. This situation is further compounded by the fact that the bulk of TE sequences present in any given genome are the evolutionary remnants of ancestral insertions. Therefore, additional methods are required to characterize comprehensively the TE landscape of non-reference genomes.

Here, we present a bioinformatic method that enables the efficient detection of TE insertion polymorphisms (TIPs) between genomes using paired-end short-reads data produced for whole-genome resequencing. This method combines an improved version of the SPLITREADER^1^ and TEPID^2^ pipelines to call the presence of non-reference TE insertions and the absence of reference TE annotations in resequenced genomes, respectively (see flowchart, Fig. 1). We apply our method to the detection of TIPs between the reference genome and ten non-reference genomes of *A. thaliana* and show that it detects TIPs with high specificity and sensitivity by comparing its performance to that of TE-capture (*Quadrana et al. 2020 Methods in Molecular Biology*).

**Figure 1.**
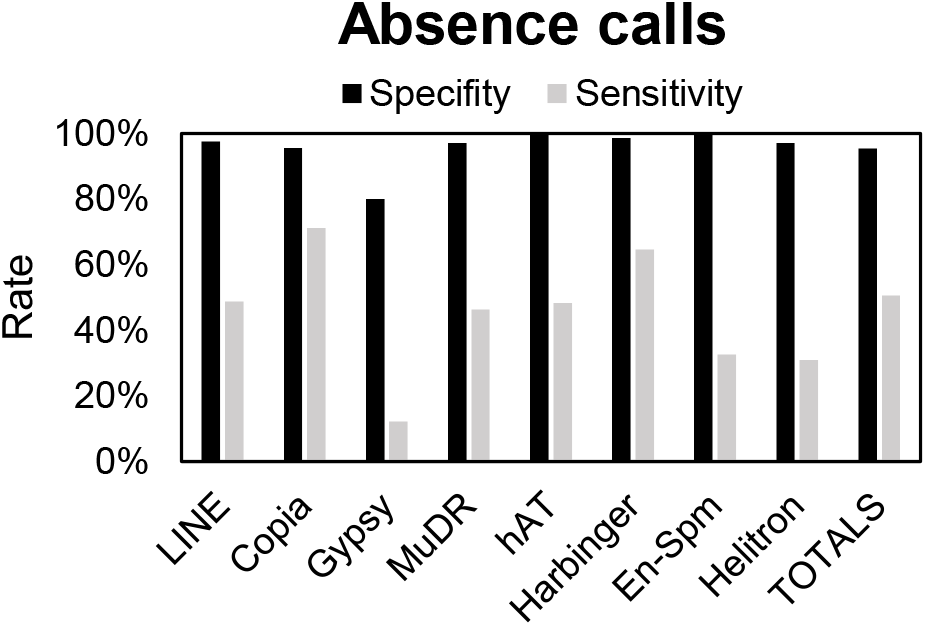
Flowchart of the SPLITREADER and TEPID pipelines used for calling non-reference TE insertion polymorphisms.

## 2 Materials

### 2.1 Software

#### 2.1.1 SPLITREADER

The updated SPLITREADER source code, split in two parts (SPLITREADER-beta2.5_part1.sh and SPLITREADER-beta2.5_part2.sh) as well as the 4 following processing scripts are available on (https://github.com/baduelp/public).

#### 2.1.2 TEPID

TEPID source code was obtained from http://doi.org/10.5281/zenodo.167274^2^.

#### 2.1.3 Other requirements

bowtie2 v2.3.2, available at https://github.com/BenLangmead/bowtie2^3^.

bedtools v2.27.1, available at https://github.com/arq5x/bedtools2^4^.

bam-readcount v0.8.0 available from https://github.com/genome/bam-readcount.

samtools v1.4.1 http://www.htslib.org/download/^5^.

Picard toolkit 2019, Broad Institute, GitHub Repository: http://broadinstitute.github.io/picard/.

### 2.2 Datasets

#### 2.2.1 Whole-genome sequencing data

Paired-ends short-read whole-genome sequencing data was obtained for 10 non-reference accessions (Table 1) from the Arabidopsis 1001 Genomes project^6^ (1001genomes.org) and for which TE-capture data was also available^1^ (*Quadrana et al. 2020 Methods in Molecular Biology*).

**Table 1.** List of 10 non-reference accessions with TE-capture data

| 1001g ID | Common name |
| --- | --- |
| 7025 | Bl-1 |
| 7343 | Sp-0 |
| 7349 | Ta-0 |
| 8264 | Bla-1 |
| 7347 | Stw-0 |
| 6911 | Cvi-0 |
| 6929 | Kondara |
| 7177 | Jm-0 |
| 7160 | Gre-0 |

#### 2.2.2 Reference genome sequence and TE annotation

Reference genome files required for the pipeline include:

- a fasta file with the reference genome sequence ($genome.fa, here TAIR10)
- a tab-delimited file with the list of TE superfamilies in the 1^st^ column and the lengths of the target-site duplication (TSD) they generate upon insertion in the 2^nd^ column (superfamily_TSD.txt)
- an annotation of TE sequences in the reference genome ($TE_annotation.gff, here TAIR10_Quesneville_GFF3)
- a list of the names of the TE families annotated in the reference genome (TE_list.txt)
- a fasta file ($TE_library.fa) with the sequences of all full-length annotated reference TEs as well as the consensus sequences for each TE family in case of degeneracy of the TE sequences present within the reference genome (e.g. *TAG1* see below).
- a tab-delimited file with the name of each TE family in the 1st column and the name of the respective superfamily in the 2nd column (TEfamily-superfamily.txt).

## 3 Methods

### 3.1 Calling non-reference TE presence variants

Given that most TIPs are expected to be rare variants, the proportion of TE presence alleles captured by the reference genome decreases exponentially with the number of genomes analyzed, each additional genome contributing to the pool of non-reference presence variants, and also to a smaller extent to the pool of reference absence variants.

#### 3.1.1 Alignment on the reference genome sequence

Illumina paired-end short reads were mapped end-to-end onto the TAIR10 reference genome sequence using Bowtie2 (using the arguments --mp 13 --rdg 8,5 --rfg 8,5 --very-sensitive) and PCR-duplicates were removed using Picard.

#### 3.1.2 Calling non-reference TE presence variants from split-and discordant reads

The detection of non-reference TE presence variants was performed by implementing an updated version of the SPLITREADER pipeline^1^. SPLITREADER has four steps: (i) extraction of reads mapping discordantly or not at all to the reference genome; (ii) mapping to a collection of reference TE sequences ($TE_library) and selection of the reads aligning partially or discordantly; (iii) re-mapping selected reads to the reference genome sequence; (iv) identification of cluster of split-reads that reveal target site duplications (TSDs).

Briefly, for each Arabidopsis accession we retrieved reads that do not map to the TAIR10 reference genome sequence (containing the SAM flag 4) or that map discordantly (paired-reads mapping to different chromosomes or to positions separated by more than 10 times the average library size). These reads were then aligned (using Bowtie2 in --local mode to allow for soft clip alignments) to a collection of 5’ and 3’ TE sequence extremities (300bp) obtained from the reference genome. In particular, for each TE family we extracted the sequence of all reference TE copies that are longer than 80% of the size of the consensus TE reported in Repbase Update7. In the case of *ARNOLDY2*, *ATCOPIA62*, *ATCOPIA95*, *TA12*, *TAG1* families, which do not contain copies with intact extremities in the reference genome, we used the TE sequence reported in Repbase Update. Next, we selected all reads mapping to a TE sequence either partially (≥20nt) or fully but with an unmapped mate. These reads were re-mapped to the TAIR10 reference genome sequence (using Bowtie2 in --local mode to allow for soft clip alignments). Read clusters composed of at least two reads mapping in the right orientation (i.e. at least one discordant read in the “+” orientation upstream of discordant read in the “−” orientation or one 3’ soft-clipped read upstream of a 5’ softclipped read, or any combination of the cases described above) were taken to indicate the presence of a *bona fide* non-reference TE presence variant.

1. Extract reads mapping discordantly or not at all to the reference genome, re-map on the $TE_library and selection of the reads aligning partially or discordantly (steps (i) and (ii)): SPLITREADER-beta2.5_part1.sh $bam_filename part1 $bam_directory $bam_extension $cohort_name $workspace_directory $TE_library
2. Re-mapping selected reads to the reference genome sequence and identification of cluster of split-reads (steps (iii) and (iv)): SPLITREADER-beta2.5_part2.sh $bam_filename $cohort_name $workspace_directory/path/to/reference/$genome $TE_annotation

#### 3.1.3 Intersecting and filtering non-reference TE presence variants by positive coverage

Putative non-reference TE presence variants sites called across each genome are then intersected and merged by TE family to define genomic regions where presence variant calls overlap. These regions are then refined and eventually split into several non-reference TE presence variants based on split-reads when available as these give precise information (at a +/− 1bp resolution) on the boundaries of non-reference TE presence variants. If split-reads are not available, some leniency (+/− 10bp) is allowed to take into account the lower precision of discordant reads in defining presence variant borders.

1. Sort (sortBed) SPLITREADER output by genome.
2. Intersect (multiIntersectBed) and sort putative presence variants by TE family ($TE_family) across genomes ($SampleNames) of the cohort ($cohort_name)
3. Merge (mergeBed) consecutive intervals of resulting file. (.sort.bed) to define overlapping intervals (.mrg.bed).
4. Refine intervals by DP filter with Filter_insertions_splitreader.pl $cohort_name $depth $project_directory $TE_family/path/to/SPLITREADER/output/by/$TE_family $TSDthresh $noTSDthresh ${SampleNames[*]} (Fig. 2):

a. Associate each merged interval (.mrg.bed) with the minimal interval it contains where most presence variants are called (.sort.bed). Set interval boundaries as flexible: $hard_start=0 and $hard_stop=0.
b. For each putative variant called in each genome, find the corresponding merged interval. Check for discrepancy between the boundaries defined in the individual genome and the boundaries of the associated interval.

i. If the individual variant and the associated interval match on both sides (+/− 1bp), add the split-reads defining the individual variant to the support of the interval boundaries ($hard_start+=$split_left and $hard_stop+=$split_right).
ii. If the individual variant and the associated interval do not match on both sides:

1. If the individual variant is defined on both sides by split-reads and the boundaries of the associated interval are still flexible ($hard_start=0 or $hard_stop=0) then refine the disagreeing boundary of the associated interval to match the one of the individual variant. Add the number of split-reads now defining the boundary to the boundary support of the associated interval ($hard_start+=$split_left or $hard_stop+=$split_right).
2. If the individual variant is defined on both sides by split-reads and the boundaries of the associated interval are not flexible ($hard_start>0 or $hard_stop>0) then create a new interval based on the individual variant.
c. For each genome, associate each putative variant called with the most frequent matching interval defined from the previous step.
d. Finally, keep non-reference TE presence variants where at least one carrier genome supports the call with at least 3 ($depth) reads (split and discordant combined with at least one split-read on each side) and:

i. The insertion size matches the expected TSD size +10bp ($TSDthresh).
ii. The insertion size is not longer than 50bp ($noTSDthresh) for TE families that do not generateTSD upon insertion.
e. Output for each presence variant, its refined boundaries, its support (number of carrier genomes as well as total number of split-reads supporting its boundaries across carriers), and its associated carriers.

**Figure 2.**
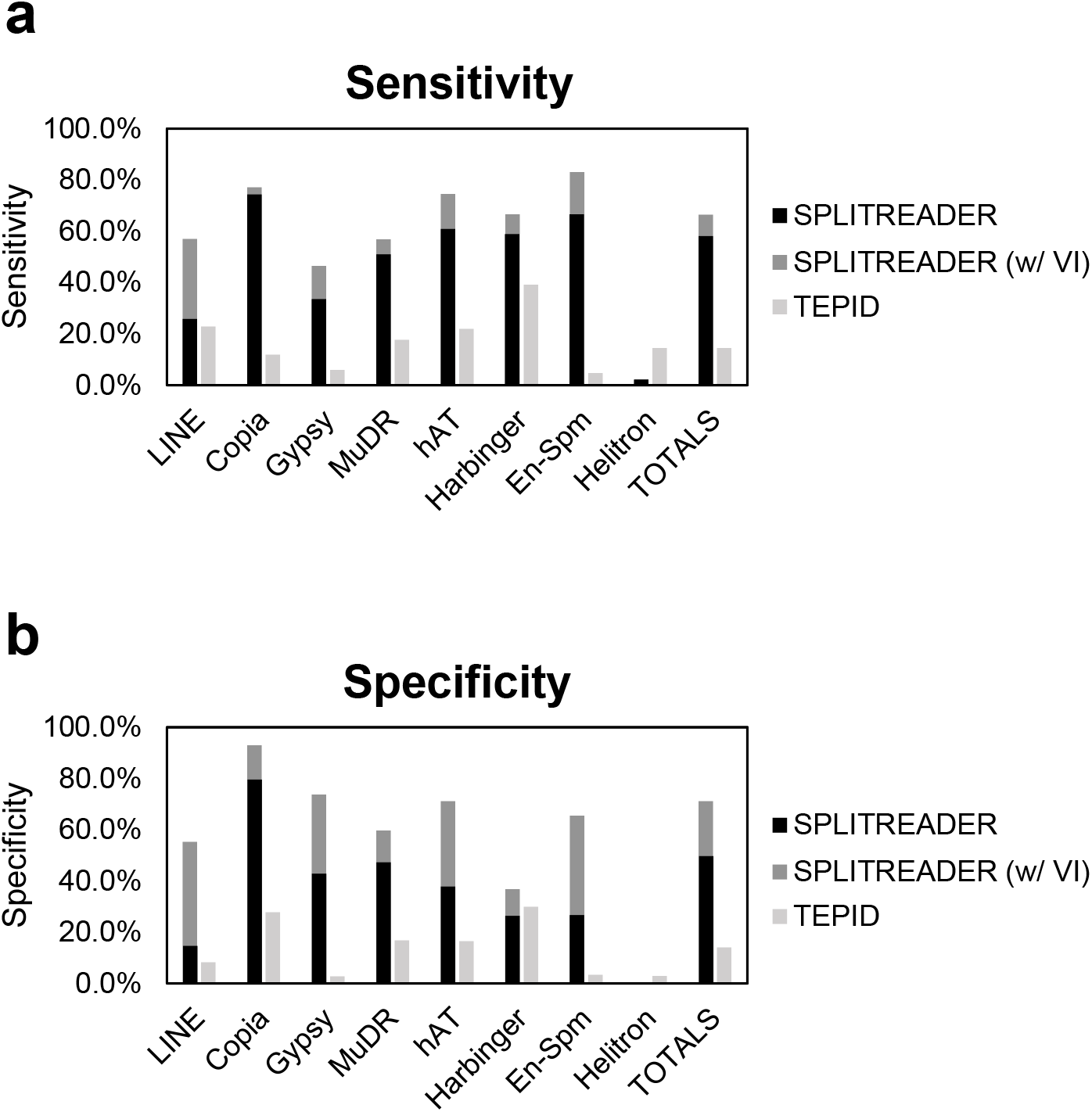
Refining intervals by DP filter. Individual SPLITREADER output intervals are indicated in grey with the number of split-reads supporting their boundaries in the circles at each extremity (when split-reads are available). Overlapping intervals are in blue with the total number of split-reads supporting their boundaries indicated at each extremity.

#### 3.1.4 Filtering non-reference TE presence variants by negative coverage drop

After the first filter (DP-filter), non-reference TE presence variants are then combined across TE-families and superfamilies to define the complete set of genomic sites for the second filter based on negative coverage (NC-filter, $filtname). Indeed, given the diploidy and high-level of homozygosity of *A. thaliana*, the negative coverage in the alignment of a carrier genome onto the reference genome should drop by at least half where non-reference TE presence variants are present (Fig. 3). Thus, the minimum coverage over the putative insertion site is compared to the average of the minimum coverage found within a similar-sized window 100bp upstream and 100bp downstream of the insertion site. In the rare cases where two different non-reference TE presence variants are called with the exact same boundaries within a given genome because of ambiguities between two closely related reference TE sequences, then only the most frequent TE presence variant is kept or that which is supported by most split-reads.

**Figure 3.**
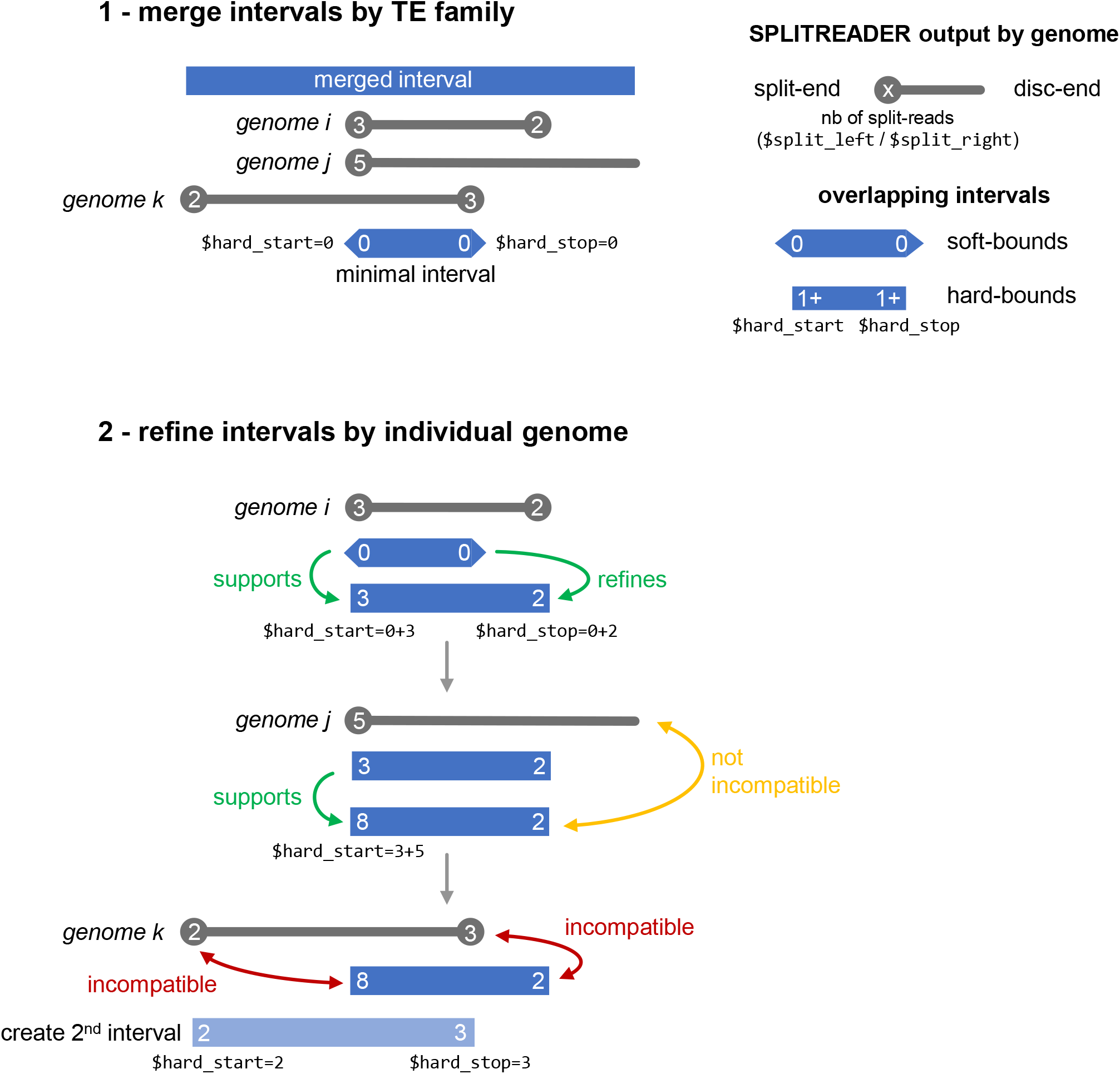
Positive and negative coverage supporting an *ATCOPIA9* insertion.

If the negative coverage does not drop by at least half or if the total number of reads over a presence variant (both supporting a presence, i.e. positive coverage, and supporting the absence by aligning on the reference genome, i.e. negative coverage) is over 100 reads, the non-reference TE presence variantis considered as a false positive for the genome in question. If an individual genome is not a carrier but the negative coverage over the presence variant is under 5 reads, it is considered as missing information or NA. Non-reference TE presence variants are kept when they fulfill all criteria listed above and have a rate of missing information below 90%.

1. Calculate negative coverage (NC) over putative insertion sites and 100bp upstream and downstream (requires Process_BAMrc_splitreader.pl) for each genome ($BAM_filename): BAM-readcount_wrapper.sh $BAM_filename BAMrc $bam_directory $bam_extension $filename $depth $cohortname $curr_dir/path/to/SPLITREADER/output/ /path/to/reference/ $genome.fasta
2. Filter presence variant calls based on negative coverage ratios (NC-filter): Filter_negative_calls_splitreader.pl $cohort_name $depth $workspace_dir $filtname ${SampleNames[*]}

### 3.2 Calling reference TE absence variants

In symmetry to the non-reference TE presence variants, a small fraction of reference TE sequences are expected to be absent from most non-reference genomes. In addition, because DNA transposons can excise, they can generate absence TE variants in non-reference genomes. To take into account these TIPss, we performed the detection of TE absence variants in the same ten non-reference genomes (Table 1) using the TEPID pipeline (Fig. 1, Stuart et al. eLife 2016). Given that TEPID performs simultaneously the calling of non-reference TE presence and reference TE absence variants, we compared its performance to that of SPLITREADER for the detection of non-reference TE insertions (see 3.3 below).

1. Map FASTQ files to the reference genome ($genome.fa): tepid-map -x /path/to/reference/$genome -y /path/to/reference/$genome.yaya_index -p $nb_cores -s $insert_size -n $filename -d /path/to/temp -o /path/to/output -1 ${filename}_1.fastq.gz -2 ${filename}_2.fastq.gz
2. Discover TE presence and absence variants by genome: tepid-discover tepid-discover --tmp /path/to/temp --split $filename.split.bam --name $filename -c $filename.bam --te /path/to/$TE_annotation.bed
3. Merge absence variants across genomes: merge_deletions.py -f deletions.list
4. Merge presence variants across genomes: merge_insertions.py -f insertions.list
5. Refine variant calls: tepid-refine -i /path/to/insertions.list.bed -d /path/to/deletions.list.bed -p $nb_cores --split $filename.split.bam--name $filename -c $bam_dir/${filename}$bam_extension.bam --te /path/to/$TE_annotation.bed -a cohort.filename.list

### 3.3 Using TE-capture to estimate the sensitivity and specificity of our TIPs calling method

In order to estimate the false-positive and false-negative rates of our TIPs calling method, we compared for ten non-reference genomes the non-reference TE presence variants detected with the SPLITREADER or TEPID pipelines or when using TE-capture ^1^ (*Quadrana et al. 2020 Methods in Molecular Biology)*. However, given that some non-reference TE presence variants can be classified as false positives when missed by TE capture (due to lower probe affinities for some TE sequences for exemple), we also performed a visual inspection of the SPLITREADER calls classified as false positives (Fig. 4).

**Figure 4.**
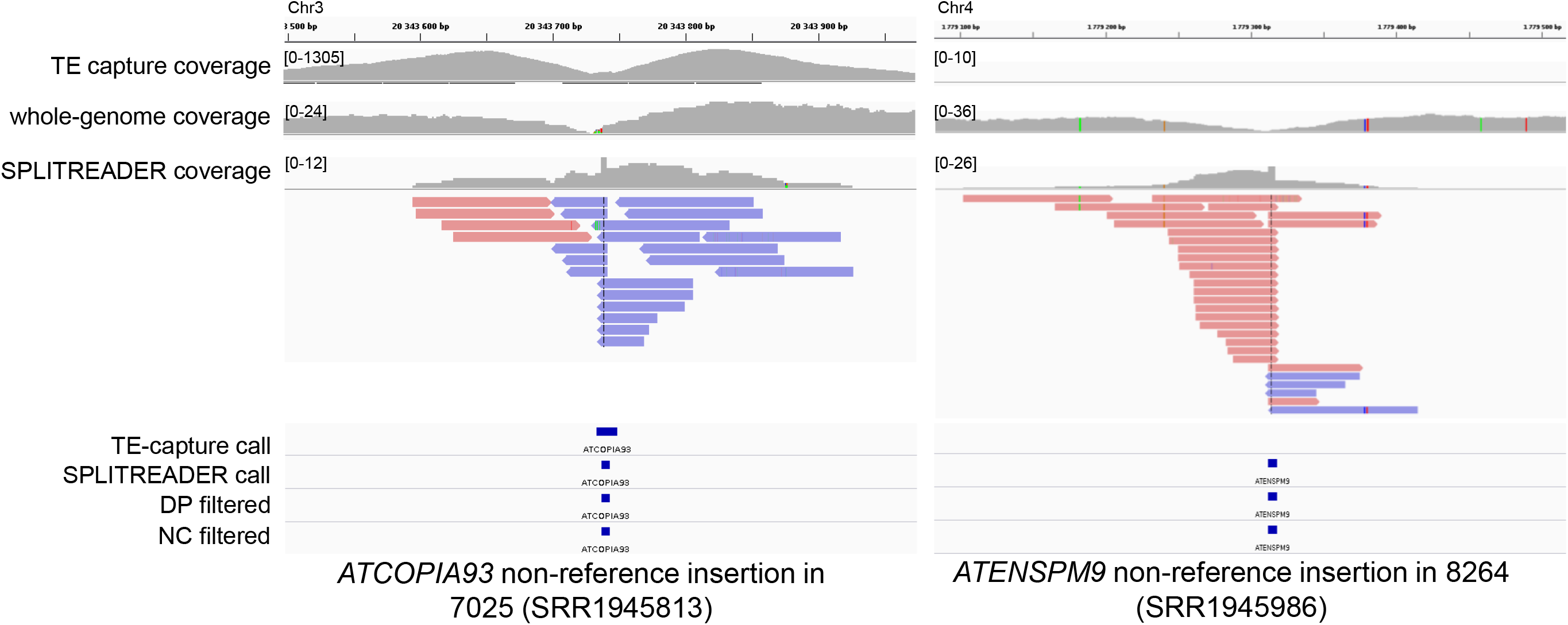
Example of two true positives called by SPLITREADER, one detected and one missed by TE-capture.

#### 3.3.1 Non-reference TE presence calls

*Helitron* DNA transposons notwithstanding, the average sensitivity of SPLITREADER reached over 66% compared to 14% for TEPID. SPLITREADER has particularly low rates of false negatives with *En-Spm* and *hAT* DNA transposons as well as with *Copia* LTR - retroelements (Fig. 5). SPLITREADER specificity was also overall very high, reaching on average 77% and 93% for *Copia* LTR-retroelements. Specificity was lowest for *LINE* retroelements (55%) and *Harbinger* DNA transposons (37%). In contrast, TEPID had low specificity overall, reaching only 14% on average.

**Figure 5.**
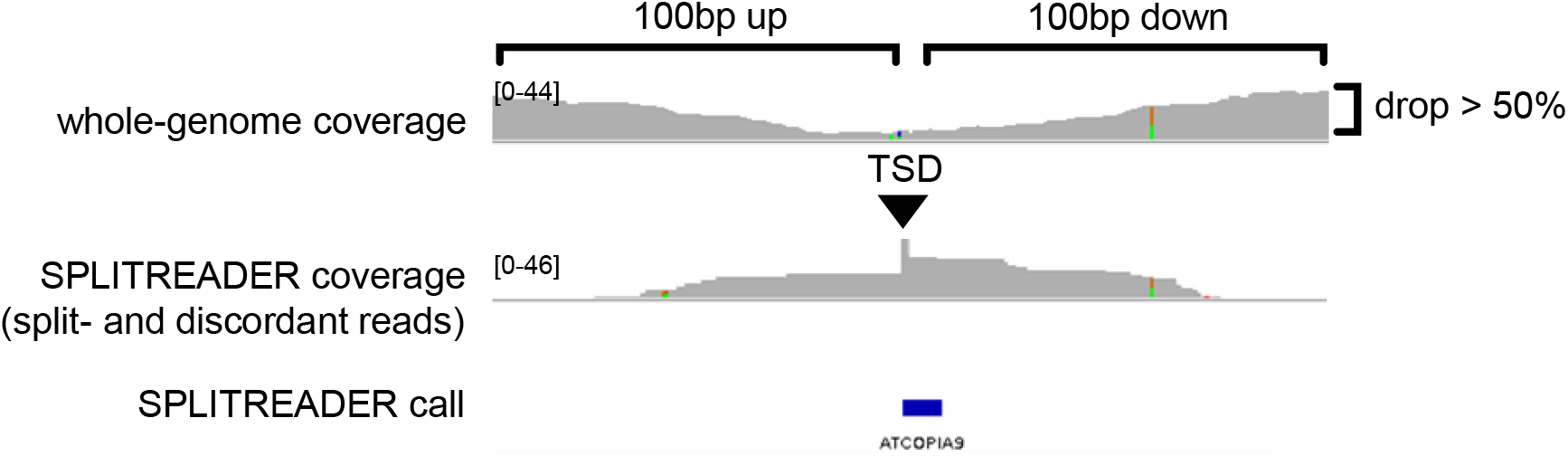
(a) Sensitivity and (b) specificity of non-reference TE presence variants calling by SPLITREADER (with or without visual inspection, VI, of false positives) and TEPID pipelines. Total rates are calculated excluding *Helitron*.

#### 3.3.2 Reference TE absence calls

Using high-coverage TE-capture from the reference genome we were also able to evaluate the sensitivity and specificity of reference TE absence calls produced by TEPID. Indeed, a reference TE absent in a non-reference genome should result in a very low sequence coverage across its annotated boundaries in the TE-capture data. Thus, when highly covered in TE-capture from the reference genome, we considered as true absence variants in TE capture from non-reference genomes the annotated TE sequences with low sequence coverage (<40 reads) within 10bp upstream and downstream outside of its boundaries but high sequence coverage (reaching >40 reads) within its boundaries. From there we estimated false-positive and false-negative rates (Fig. 6).

**Figure 6.**
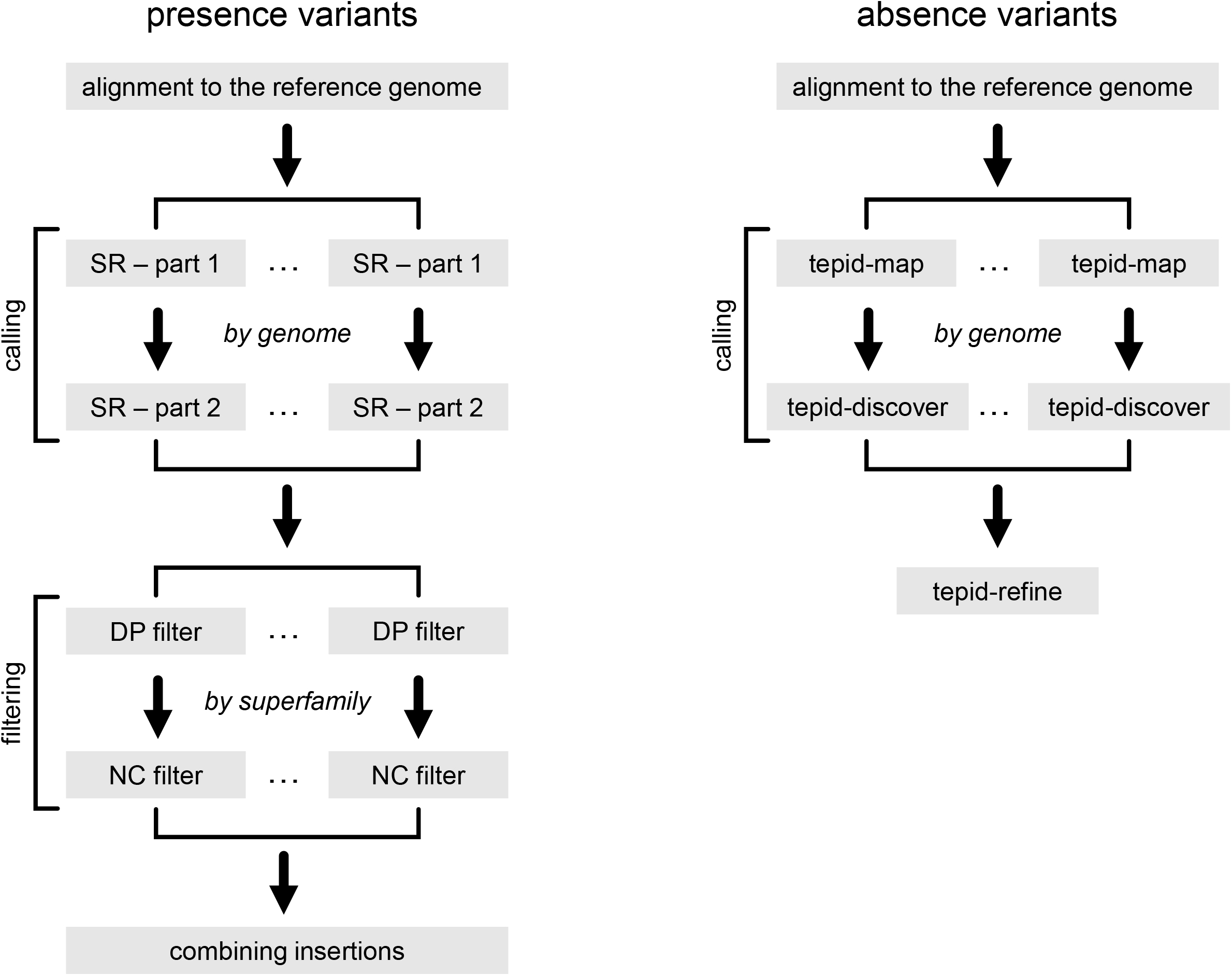
Sensitivity and specificity of reference TE absence variants called by TEPID pipeline.

Across all TE superfamilies including *Helitron*, specificity of reference TE absence calling by TEPID was very high, reaching on average 95%, with a minimum of 80% for *Gypsy* retroelements. Sensitivity was lower however, reaching 50% on average, with a maximum of 70% for *Copia* retroelements and a minimum of 12% for *Gypsy* retroelements.

## 4. Conclusion

We show that by combining TEPID with an updated version of SPLITREADER, TIPs can be detected readily across multiple genomes with high-sensitivity and high-specificity using short-read sequencing data. Although we can expect that the advent of long-read sequencing will result in a rapid increase of the number of fully assembled genome sequences, thus allowing to detect TIPs and their exact sequence comprehensively, short reads will likely remain the most common approach for population genomic studies. We therefore anticipate that our method for TIP detection will find broad applications.

## 5. Acknowledgments

We thank members of the Colot group for discussions. P.B. was the recipient of a postdoctoral fellowship (code SPF20170938626) from the Fondation pour la Recherche Médicale (FRM). Support was from the Agence National de la Recherche (ANR-09-BLAN-0237, the Investissements d’Avenir ANR-10-LABX-54 MEMO LIFE, ANR-11-IDEX-0001-02 PSL* Research University to V.C) and the Centre National de la Recherche Scientifique (MOMENTUM program, to L.Q.).

